# Modeling place cells and grid cells in multi-compartment environments: hippocampal-entorhinal loop as a multisensory integration circuit

**DOI:** 10.1101/602235

**Authors:** Tianyi Li, Angelo Arleo, Denis Sheynikhovich

## Abstract

Hippocampal place cells and entorhinal grid cells are thought to form a representation of space by integrating internal and external sensory cues. Experimental studies show that different subsets of place cells are controlled by vision, self-motion or a combination of both. Moreover, recent studies in environments with a high degree of visual aliasing suggest that a continuous interaction between place cells and grid cells can result in a deformation of hexagonal grids or in a progressive loss of visual cue control. The computational nature of such a bidirectional interaction remains unclear. In this work we present a neural network model of a dynamic loop between place cells and grid cells. The model is tested in two recent experimental paradigms involving double-room environments that provide conflicting evidence about visual cue control over self-motion-based spatial codes. Analysis of the model behavior in the two experiments suggests that the strength of hippocampal-entorhinal dynamical loop is the key parameter governing differential cue control in multi-compartment environments. Construction of spatial representations in visually identical environments requires weak visual cue control, while synaptic plasticity is regulated by the mismatch between visual- and self-motion representations. More gener-ally our results suggest a functional segregation between plastic and dynamic processes in hippocampal processing.

## 1. Introduction

It has long been accepted that spatial navigation depends crucially on a combination of visual and self-motion input (O’Keefe and Nadel, 1978). Since the seminal work of O’Keefe and Dostrovsky (1971), a neural locus of this combination is thought to be the place cell network in the CA1-CA3 subfields of the hippocampus proper (O’Keefe and Speakman, 1987, Muller and Kubie, 1987, Knierim et al., 1998, Jayakumar et al., 2018), with different subsets of place cells sensitive to self-motion cues, to visual cues or, more often, to a combination of them (Markus et al., 1994, Chen et al., 2013, Fattahi et al., 2018). A more recent discovery of grid cells in the medial entorhinal cortex led to the suggestion that the grid-cell network provides a self-motion-based representation of location that is combined with other sensory information on the level of place cells (Fyhn et al., 2004, McNaughton et al., 2006, Hayman and Jeffery, 2008, Cheng and Frank, 2011). The grid-cell representation is itself vision-dependent, since various properties of grid cells are affected by changes in visual features of the environment (Hafting et al., 2005, Krupic et al., 2015). Combined with the evidence showing that coherent changes in place-cell and grid-cell representations occur during environment deformation and cue manipulation, these data suggest a bidirectional interaction between these representations at the neural level (Fyhn et al., 2007). While this bidirectional link is always present in normal conditions, it may not be necessary for place cell activities, as shown in a number of lesion experiments (Sasaki et al., 2015, Schlesiger et al., 2018).

The nature of the dynamic interaction between visual and self-motion cues on the level of grid cells has recently been tested in two experiments: in a merged room, formed by removal of a wall separating two visually similar environments (Wernle et al., 2018), and during exploration of an environment consisting of two identical rooms connected by a corridor (Carpenter et al., 2015). Results of the first experiment have shown that firing patterns of grid cells were anchored by local sensory cues near environmental boundaries, while they underwent a continuous deformation far from the boundaries in the merged room, suggesting a strong control of local visual cues over grid-cell representation (Wernle et al., 2018). Results of the second experiment indicated in contrast that during learning in a double-room environment grid cells progressively formed a global self-motion-based representation disregarding previously learned local cues (Carpenter et al., 2015).

Existing models of the entorhinal-hippocampal system are mostly based on the feed-forward input from grid cells to place cells, with an additional possibility to reset grid-field map upon the entry to a novel environment (Solstad et al., 2006, O’Keefe and Burgess, 2005, Blair et al., 2008, Sheynikhovich et al., 2009, Pilly and Grossberg, 2012), or focus on the feed-forward input from place cells to grid cells (Bonnevie et al., 2013). In addition to be at difficulty at explaining the above results on dynamic interactions between visual and self-motion cues, they are also not consistent with data showing that hippocampal spatial representations remain spatially tuned after MEC inactivation (Brun et al., 2008, Rueckemann et al., 2016) and that in pre-weanling rat pups, place fields can exist before the emergence of the grid cell network (Muessig et al., 2015). Moreover, disruption of grid cell spatial periodicity in adult rats does not alter preexisting place fields nor prevent the emergence of place fields in novel environments (Koenig et al., 2011, Brandon et al., 2014).

In this paper we propose a model of continuous dynamic loop-like interaction between grid cells and place cells, in which the main functional parameter is the feedback strength in the loop. We show that the model is able to explain the pattern of grid-cell adaptation in the two experiments by assuming a progressive decrease of visual control over self motion, and a plasticity mechanism regulated by allothetic and idiothetic cue mismatch over a long time scale.

## 2. Model

This section presents main neuronal populations in the model and their interactions. Further technical details and model parameters are given in the Appendix.

The rat is modeled by a panoramic visual camera that is moving in an environment along quasi-random trajectories resembling those of a real rat (Fig. 2A, top). The orientation of the camera corresponds to the head orientation of the model animal. The constant speed of the modeled rat is set to 10 cm/s, and sampling of sensory input occurs at frequency 10 Hz, roughly representing hippocampal theta update cycles. The modeled rat receives two types of sensory input (Fig. 1). First, self-motion input to the model is represented by angular and translational movement velocities integrated by grid cells in the medial entorhinal cortex (mEC) to provide self-motion representation of location, as proposed earlier (McNaughton et al., 2006). Competitive self-organization of grid cell output occurs downstream from the entorhinal cortex in the dentate gyrus (DG) - CA3 circuit and gives rise to a self-motion-based representation of location, encoded by *motion-based place cells* (MPC). We did not include a specific neuronal population to model DG (de Almeida et al., 2009a). Instead, we implemented competitive learning directly on mEC inputs to CA3. Second, visual input is represented by responses of a two-dimensional retina-like grid of orientation-sensitive Gabor filters, applied to input camera images at each time step. For instance, in featureless rectangular rooms used in most of the simulations below, the only features present in the input images are the outlines of the environment walls (Fig. 2A, bottom). Importantly, the ‘retinal’ responses are assumed to be aligned with an allocentric directional frame further along the dorsal visual pathway (not modeled), the directional frame being set by head direction cells (Byrne et al., 2007, Sheynikhovich et al., 2009, Bicanski and Burgess, 2018). That is, visual input to the model at each spatial location is independent on the head direction that the model rat has upon arriving at that location. The visual input aligned with an allocentric directional frame is assumed to be encoded in the inputs to the hippocampal formation from the lateral entorhinal cortex (lEC). Competitive self-organization of these inputs results in a purely vision-based representation of location, encoded by a population of *visual place cells* (VPCs). Both MPCs and VPCs project to CA1 cells that form a conjunctive representation of location in *conjunctive place cells* (CPCs). The principal novelty of the model is that CPCs in CA1 project back to the entorhinal grid cells and thus form a recurrent loop, reflecting the anatomy of entorhinal-hippocampal connections (Iijima et al., 1996).

**Figure 1:**
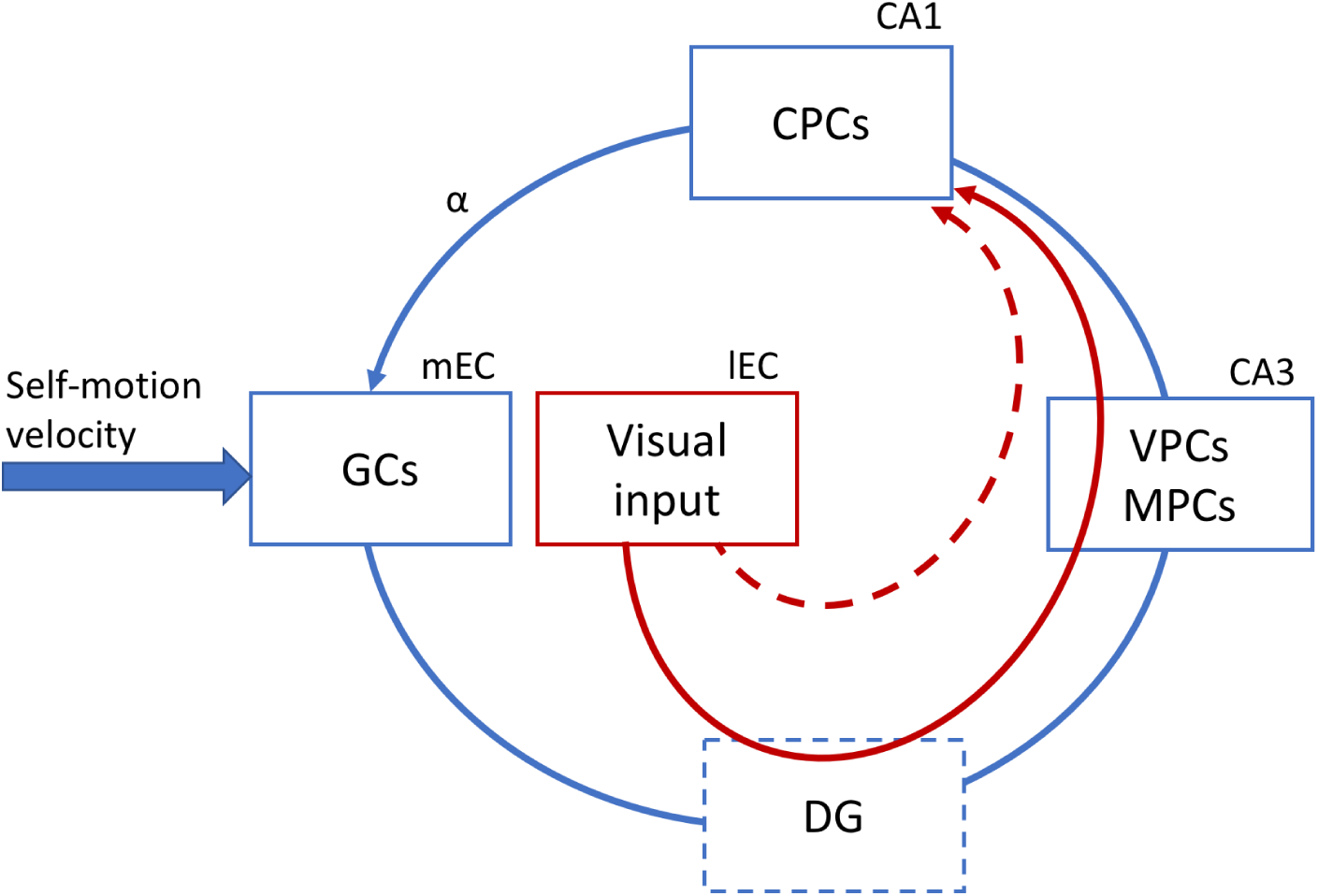
_Schematic representation of the model. Self-motion input is integrated in the grid cell populations of the medial EC, and via competitive interactions results in a self-motion-driven space representation in CA3 (encoded by the MPC population). Visual input, coming via the lEC, results in a purely vision-based representation in CA3, encoded by the VPC population. Both MPCs and VPCs project to CA1 where the conjunctive representation of location is encoded in the CPC population. The projection from CPCs in CA1 back to the mEC closes the dynamic hippocampal processing loop and the strength of this projection is determined by the parameter_ *_α_*_. The full arrows represent the information flow in the network. The dashed arrow represents an alternative way to model visual input processing. The DG is not modeled._

**Figure 2:**
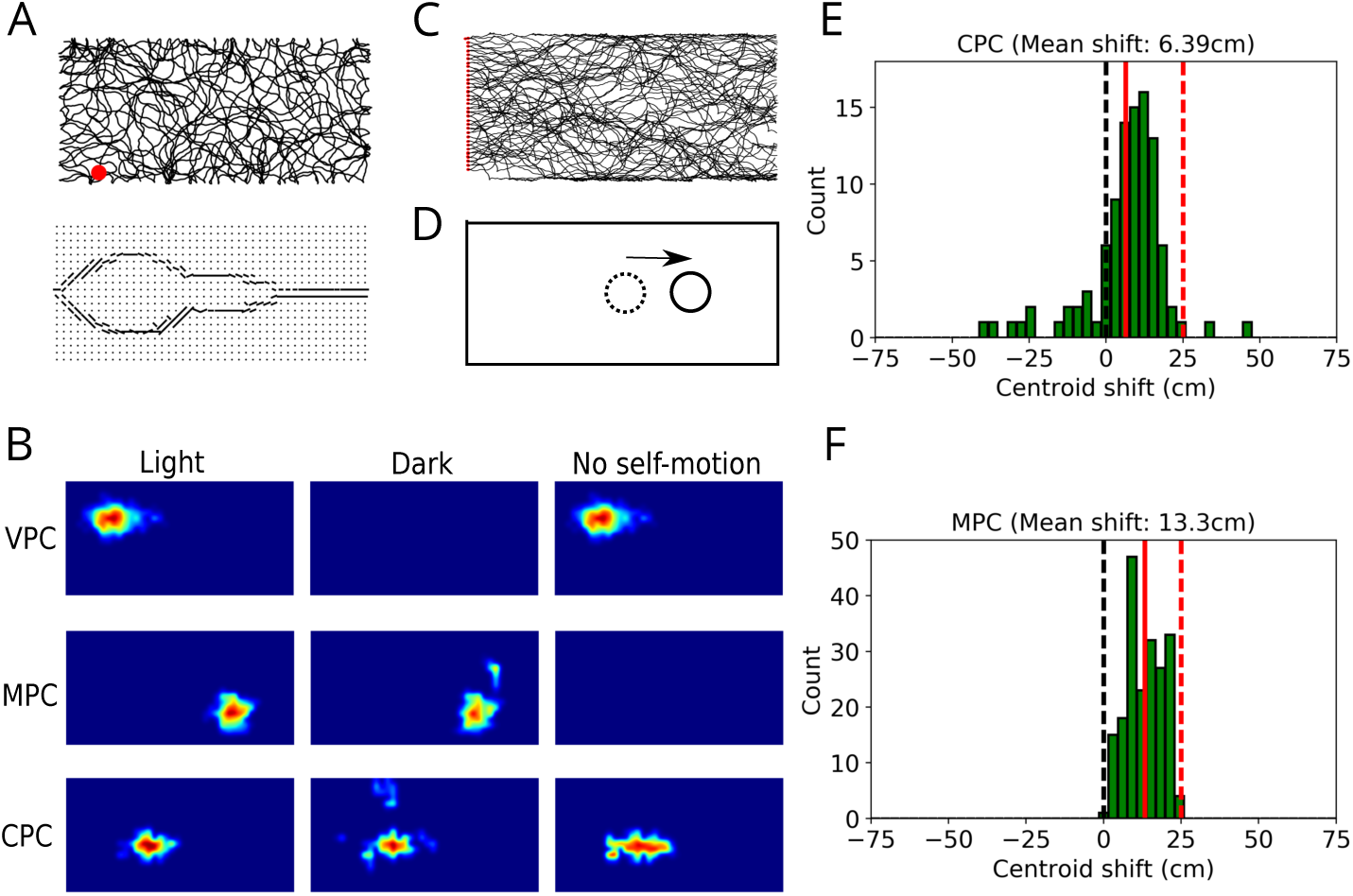
Multisensory integration in modeled place cells. A. An example of the trajectory of the modeled animal in a rectangular environment (top) and the visual input to the model (bottom) from the location marked by the red dot. In the bottom plot, the dots represent the grid of Gabor filters, and lines represent the orientations of most active filters. Visual input at each location is independent from head direction. B. Firing fields of VPCs (top row), MPCs (middle row) and CPCs (bottom row) in simulated ‘light’ condition (left column), ‘dark’ condition (middle column) and passive translation (right column). C. Trajectories of model animal crossing the rectangular environment from left to right. The red dots denote the starting positions. D. When the model rat crosses the environment from left to right, self-motion position estimate (dotted circle) is behind the visual position estimate (full circle) in the conditions of decreased speed gain, leading to a forward-shift of receptive fields. E,F. Forward-shift of receptive fields in the population of CPCs (top) and MPCs (bottom). Full red lines represent the mean shift in the population. Dashed red lines represent the shift due to purely self-motion input.

### Integration of visual and self-motion input by grid cells

The self-motion input is processed by 5 identical neuronal populations representing distinct grid cell populations in the dorsal mEC (Hafting et al., 2005). Each grid cell population can be represented as a two-dimensional sheet of neurons equipped with attractor dynamics on a twisted-torus topology, as has been proposed in earlier models (Guanella et al., 2007, Sheynikhovich et al., 2009, Burak and Fiete, 2009). The position of an attractor state (or *activity packet*) in each grid cell population is updated based on the self-motion velocity vector. This is implemented by the modulation of recurrent connection weights between grid cells according to the model rat rotation and displacement, such that the activity bump moves across the neural sheet according to the rat movements in space (Guanella et al., 2007). The only difference between grid-cell populations is that the speed of movement of the activity bumps across the neural sheet is specific for each population, resulting in population-specific distance between neighbouring grid fields and field size (Hafting et al., 2005). As long as each location in an environment corresponds to a distinct combination of positions of the activity packets, population activity of all grid cells encodes the current position of the animal in the environment (Burak and Fiete, 2009). The exact implementation of the attractor mechanism governing grid-cell network dynamics is not essential for the model to work.

In addition to the recurrent input from grid cells in the same population, each grid cell receives input from the CPC population which represent conjunctive visual and self-motion representation (described in detail later), and the relative strength of these two inputs is controlled by the parameter *α*. At a relatively high value of this parameter, grid-cell attractor dynamics in each layer is strongly influenced by the hippocampal input, leading to an overall stronger effect of visual information. At a low value of *α*, the grid-cell dynamics is governed almost exclusively by self-motion input.

Thus, the total synaptic input to a grid cell *i* at time *t* is (omitting grid cell population index for clarity)

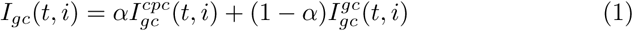

where the external input from CPC and recurrent inputs from other grid cells are determined by

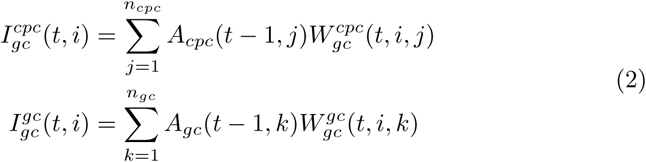

Here, *A_cpc_*(*t, j*) is the activity of *j*-th CPC at time *t* (described below) and *A_gc_*(*t, k*) = *I_gc_*(*t, k*) is the activity of *k*-th grid cell (we use linear activation function for grid cells).

Feedforward synaptic connections from CPCs are initialized by random values and updated during learning according to a standard Hebbian learning scheme:

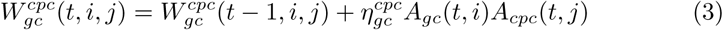

followed by explicit normalization ensuring that the norm of the synaptic weight vector of each cell is unity (a neurally plausible implementation of the normalization step can be implemented by a change in the learning rule (Oja, 1982)).

Recurrent synaptic connections between grid cells are constructed such as to ensure attractor dynamics, modulated by velocity vector (Guanella et al., 2007). More specifically, the connection weights between cells *i* and *j* is a Gaussian function of the distance between these cells in the neural sheet. This connection weight is modulated by the self-motion velocity vector, such that the activity bump moves across the neural sheet according to the direction and norm of the velocity vector, with a proportionality constant that is grid-cell population specific. These proportionality constants were tuned such that the grid spacing across different grid cell populations were between 42 cm and 172 cm. Grid-cell firing patterns were oriented 7.5° with respect to one of the walls of an experienced experimental enclosure (Krupic et al., 2015).

### Encoding of visual and self-motion input by place cells

As mentioned above, the model includes three distinct populations of place cells (Fig. 1). First, VPCs directly integrate allocentric visual inputs, presumably coming from lEC and project further to CA1. We putatively assign VPC population to CA3 where a competitive mechanism based on recurrent feedback can result in self-organization of visual inputs, the resulting spatial code further transmitted to to CA1. The model of this pathway is based on the evidence that stable spatial representations were observed in CA1 after complete lesions of the mEC containing grid cells (Brandon et al., 2014, Schlesiger et al., 2018). Second, MPCs directly integrate input from grid cells and in the absence of visual inputs the activity of these cells represents purely self-motion-based representation of location. These cells represent CA3 place cells, acquiring their spatial selectivity via a competitive mechanism based on mEC inputs (de Almeida et al., 2009a). Third, CPCs that model CA1 pyramidal cells, combine visual and self-motion inputs coming from VPC and MPC populations, respectively. Crucially, CPCs project back to the grid cell populations, modeling anatomical projections from CA1 back to the entorhinal cortex forming a loop (Iijima et al., 1996, Slomianka et al., 2011) and controlled by the parameter *α* as described above.

#### Vision-based place cells

VPCs acquire their spatial selectivity as a result of unsupervised competitive learning implemented directly on allocentric visual inputs, represented by Gabor filter activities aligned to an allocentric directional frame (see Appendix). As a result of learning, different cells become sensitive to constellations of visual features observed from different locations (independently from head direction).

The total input to a VPC *i* at time *t* is given by

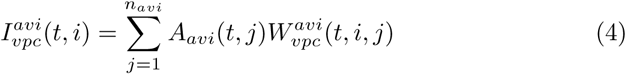

where *A_avi_*(*t, j*) is the activity of *j*-th Gabor filter aligned with the allocentric directional frame. A E%-max winner-take-all learning scheme (de Almeida et al., 2009a,b) is implemented, meaning that a small subset of maximally active cells is selected (i.e. all cells whose total input is within *E_vpc_*% of the cell with maximal input). The synaptic weight updates according to the Hebbian modification rule (Eq. 3) are implemented only for the winner cells.

#### Motion-based place cells

MPCs read out grid cell activities similarly to previously proposed models (Solstad et al., 2006, Sheynikhovich et al., 2009). More specifically, they implement the E%-max winner-take-all learning scheme identical to that of VPCs learning described above (with parameter *E_mpc_* determining the proportion of highly active cells).

#### Conjunctive place cells

Both VPCs and MPCs project to CPCs, that model CA1 pyramidal cells sensitive to both visual and self-motion cues. The total input to a conjunctive cell is:

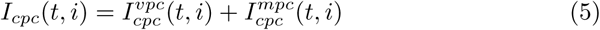

With

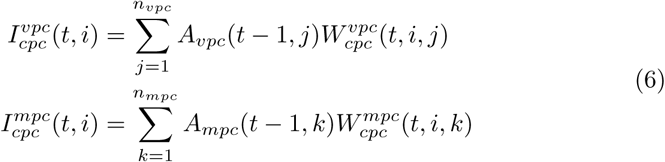

Again, a E%-max winner-take-all learning scheme is implemented in this network, but with a heterosynaptic update learning rule:

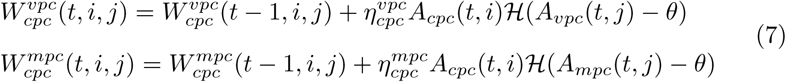

where *H*(.) is the Heaviside step function (*H*(*x*) = 0 for *x ≤* 0, and *H*(*x*) = *x* otherwise) and *θ* is the presynaptic activity threshold.

Due to the fact that MPCs, CPCs and grid cells are connected in a loop, a local activity packet in an “upstream” cell population shifts the activity packet in the “downstream” population towards the position of former. The size of the induced shift on each cycle of theta is determined by connection strengths between participating cells. In the absence of visual input, activity bumps in the three interconnected populations settle at the global stable state of the loop/attractor dynamics and hence all code for a single spatial location in the environment, which can be considered as the estimation of the animal’s location based on self-motion input. However, because of the visual input from VPCs, the loop dynamics is biased towards the visual position, encoded in the VPC population. Thus, the feedback strength in the loop determines the extent to which visual input influences place cell activities in the model.

## 3. Results

Since the early experiments testing the influence of visual and self-motion cues on place cell activity, it was clear that different subsets of place cells are controlled by these cues to different degrees, with some cells being controlled exclusively by one type of cue (Markus et al., 1994, Chen et al., 2013, Aronov and Tank, 2014, Fattahi et al., 2018). In the model we conceptualized these differences in VPC, MPC and CPC neural populations, representing purely vision-dependent, motion-dependent and multisensory place cells. Thus, when the model has learned place fields in a visually structured environment by moving quasi-randomly around a rectangular box, VPCs have place fields only in a ‘light’ condition, i.e. when the visual cues are visible. This is true even if motion-based cues are absent (Fig. 2B, top row), as in a passive transport through a virtual maze (Chen et al., 2013). Conceptually, these cells represent the ability of hippocampal circuits to form self-organized representations of location even in the absence of grid-cell input from the mEC (Hales et al., 2014, Brandon et al., 2014, Schlesiger et al., 2018). In contrast, MPCs will have place fields both in the light and dark conditions, but not during passive translation (Fig. 2B, middle row). Finally, CPCs will be active in all the three conditions since they combine both types of input (Fig. 2B, bottom row).

In contrast to VPCs that are completely independent of self-motion cues and encode stable visual features of the surrounding environment, MPCs and CPCs will be influenced by both visual and self-motion input, by virtue of their loop-like interactions through the grid cells. To test the relative influence of vision and self motion on the activity of these cells when the two types of cue provide conflicting sensory information, we decreased the gain of self-motion input to grid-cells while the model animal was crossing the environment from left to right (Fig. 2C). This decrease in gain was applied only to the horizontal component of motion, i.e. the horizontal component of the self-motion velocity vector was set to 3*/*4 of the baseline value. Such a modulation is similar to a change in the gain of ball rotation in a virtual corridor (Chen et al., 2013), but implemented in a two-dimensional environment instead of a linear track. The change in gain resulted in a shift of receptive fields of MPCs and CPCs to the right relative to their position in baseline conditions and the size of the shift is smaller than what would be predicted from purely self-motion integration (Figs. 2E,F).

To illustrate the loop dynamics in this simple example, consider the case when the model animal crosses the middle line of the environment moving from left to right (Fig. 2D). The integration of pure self-motion input over time would estimate the current position to be behind the visually estimated position due to the decrease in speed gain. This will cause a cell that normally fires at the center of the environment to shift its receptive field ahead of it. Thus, in the dark condition MPCs and CPCs have place fields shifted forward by an amount proportional to the gain factor, relative to their positions in the baseline condition (i.e. without the change in gain). However, in the light condition this self-motion-based estimation will be in conflict with visual cues that are not affected by changes in gain and represent the actual position in the environment. As a result of the dynamic loop-like interaction, at each moment of time visual cues induce a forward shift of the activity packet in the grid-cell populations towards the visually identified location, the size of the shift being controlled by the parameter *α*. Grid cells would similarly affect the MPCs, and then CPCs, closing the loop. Therefore, in the presence of conflicting cues receptive fields shift to an intermediate position between the self-motion and visual estimates (Figs. 2E,F). These results are reminiscent of those by Gothard et al. (1996), simulated in several earlier computational models (Samsonovich and McNaughton, 1997, Byrne et al., 2007, Sheynikhovich et al., 2009), and indeed the proposed mechanistic explanation is similar in this case. However, in the present model the parameter controlling the interaction between the visual and self-motion cues is cast in terms of the strength of the entorhinal-hippocampal loop.

To illustrate the same multisensory integration mechanism on the level of grid cells, we conducted another simulation in which the horizontal velocity gain was transiently decreased when the model animal crossed a specific portion of the environment (Fig. 3A). In this case of a transient cue conflict, grid patterns were locally deformed in that firing fields near the gain-decrease zone shifted to the right relative to control conditions, reflecting the sensory conflict (Figs. 3B-D). Near the borders of the environment, where the speed input was identical to the baseline conditions, grid pattern remained stable. The same effect on the level of the whole population of grid cells was quantified by the analysis of displacement vectors (Fig. 3C) and by sliding correlation maps (Fig. 3D), see Appendix and Wernle et al. (2018). These results suggest that local modifications of grid patterns can be induced by conflicting sensory representations, similarly to what has been observed in a recent experiment by Wernle et al. (2018). As mentioned in the Introduction, these results are at odds with an earlier experiment (Carpenter et al., 2015) that studied adaptation of grid-cell patterns during construction of a spatial representation in an environment consisting in two identical rooms connected by a corridor. In the following sections we simulated the results of both experiments in an attempt to explain this conflict and to understand neural mechanisms responsible for apparently different patterns of grid-cell adaptation in the two experiments.

**Figure 3:**
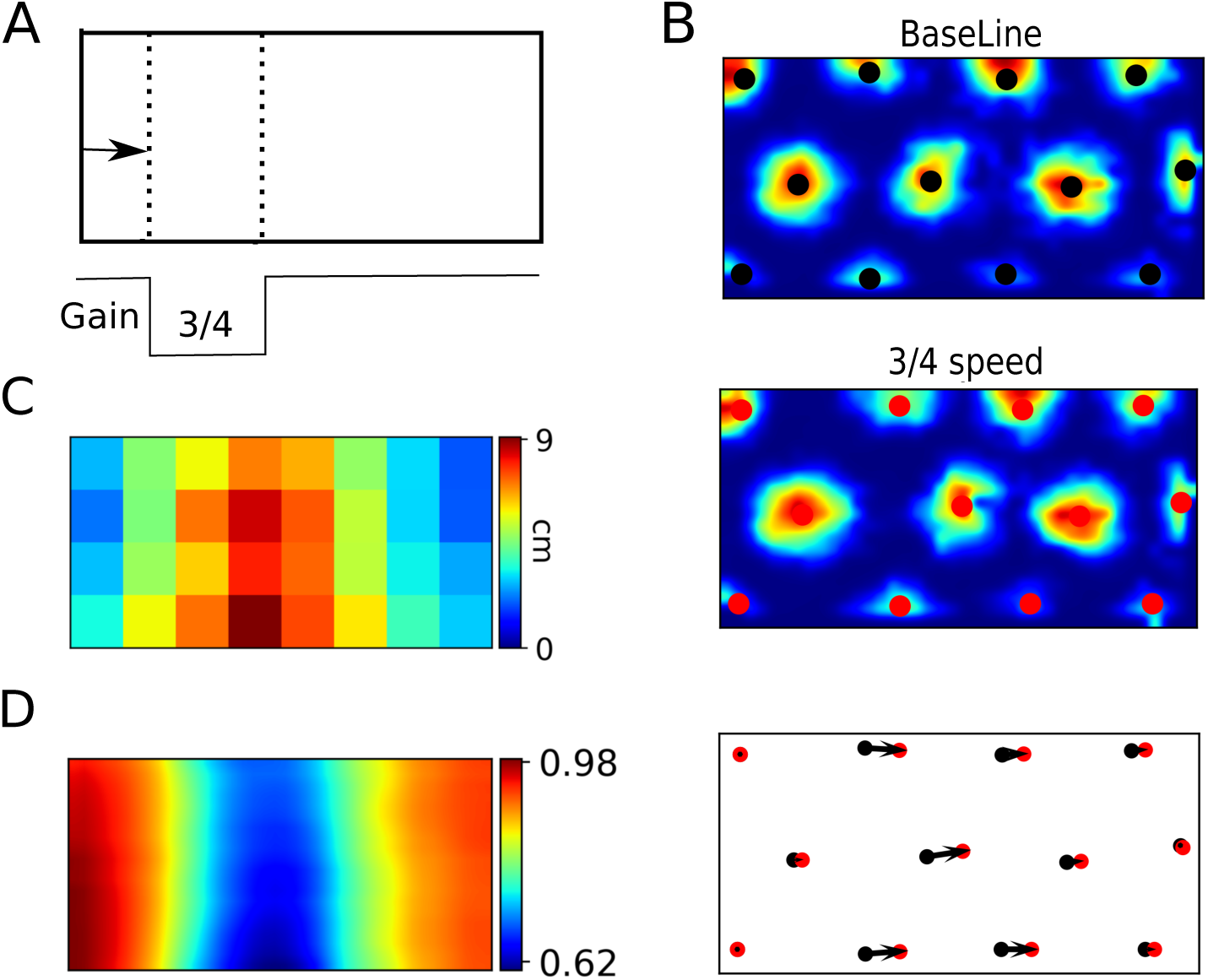
Multisensory integration in grid cells. A. The speed gain was transiently decreased to 3/4 of the normal gain when the model animal approached the portion of the environment marked by the dotted lines. B. An example of firing pattern of a grid cell in the conditions of normal speed (top) and with transiently decreased speed gain (middle). The black and red circles represent the centers of firing fields in the baseline condition and during decreased gain, respectively. The shift of firing fields is quantified by displacement vectors shown by the black arrows (bottom). C. Color map of the mean displacement vector lengths in different portions of the environment. D. Color map of mean sliding correlation over all grid cells.

### 3.1. Merged-room experiment

Wernle et al. (2018) studied the integration between visual and self-motion cues by recording grid cells in two adjacent rectangular compartments initially separated by a wall. The two compartments were inserted in a bigger environment equipped with distal visual cues. The wall was subsequently removed and grid cells were recorded while the rat foraged in the merged environment. The authors observed that at the locations far from the removed wall grid cells conserved their firing patterns, while at the locations near those previously occupied by the wall grid-cell firing fields shifted towards the removed wall so as to form a continuous quasi-hexagonal pattern. Results from the previous section suggest that the observed local deformation of the grid pattern can result from the local visual deformation caused by wall removal.

To verify that our model can reproduce these results, we recorded activities of simulated grid cells and place cells cells in experimental conditions similar to those in Wernle et al. More specifically, the model learned place fields in two virtual rooms separated by a wall (Fig. 4A). The two rooms were located inside a bigger room with distal visual cues (not shown), such that learned representations of the two rooms were different after initial exploration. After place fields were established, the wall was removed, the synaptic weights were fixed and neural activity was recorded. We observe that after wall removal, grid fields near distant walls remain fixed to the local cues, while near the former wall location they shift towards this location in the model, as in the experiment (Fig. 4B). The same phenomenon on the level of the whole population was quantified by the analysis of displacement vectors (Fig. 4C) and by sliding correlation (Fig. 5D).

**Figure 4:**
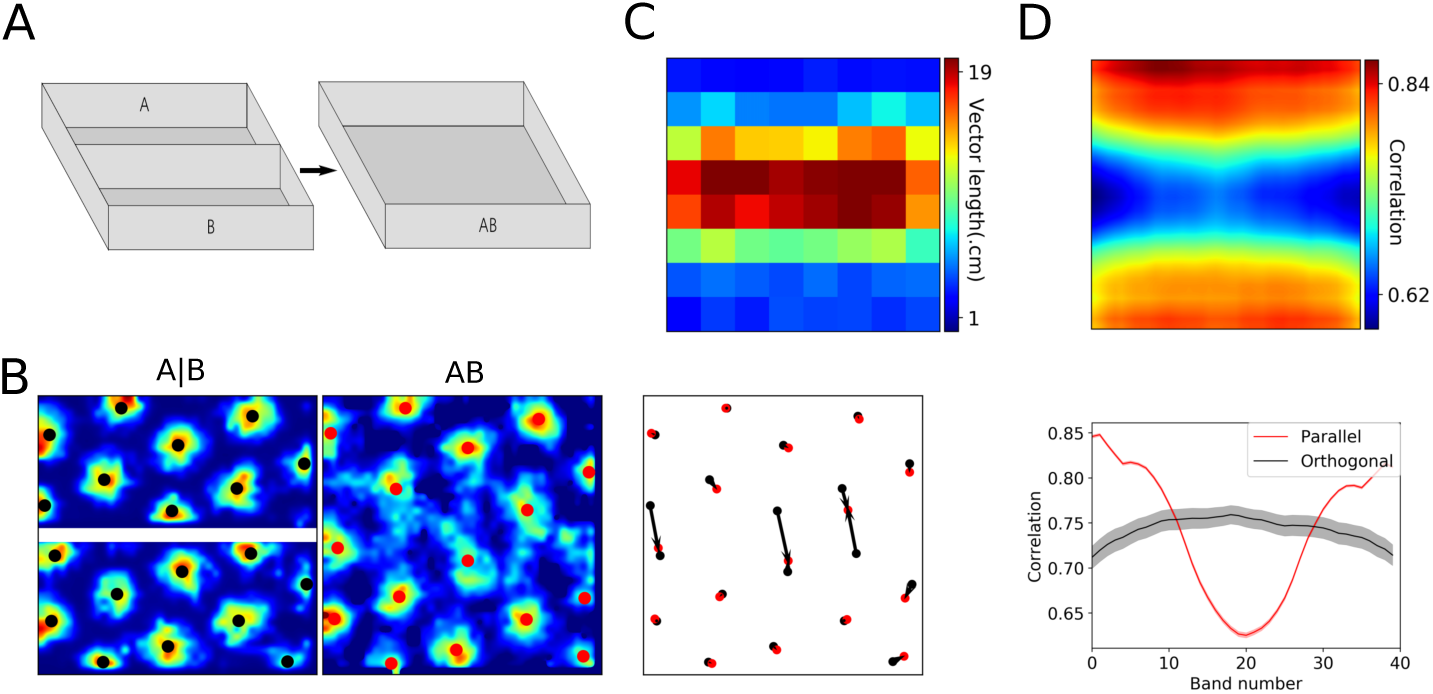
Simulation of the merged-room experiment of Wernle et al. (2018). A. The training environment with two separate rooms, referred to as room ‘A B’, and the testing environment, referred to as merged room ‘AB’. B. Firing fields of an example grid cell in the training (left) and testing (middle) environments, as well as firing-field displacement vectors calculated in the testing environment (right). C. A color map of mean vector lengths. D. Top plot: A color map representing the mean sliding correlation over all grid cells. Bottom plot: the correlation profiles at the center of the environment along two cardinal directions.

**Figure 5:**
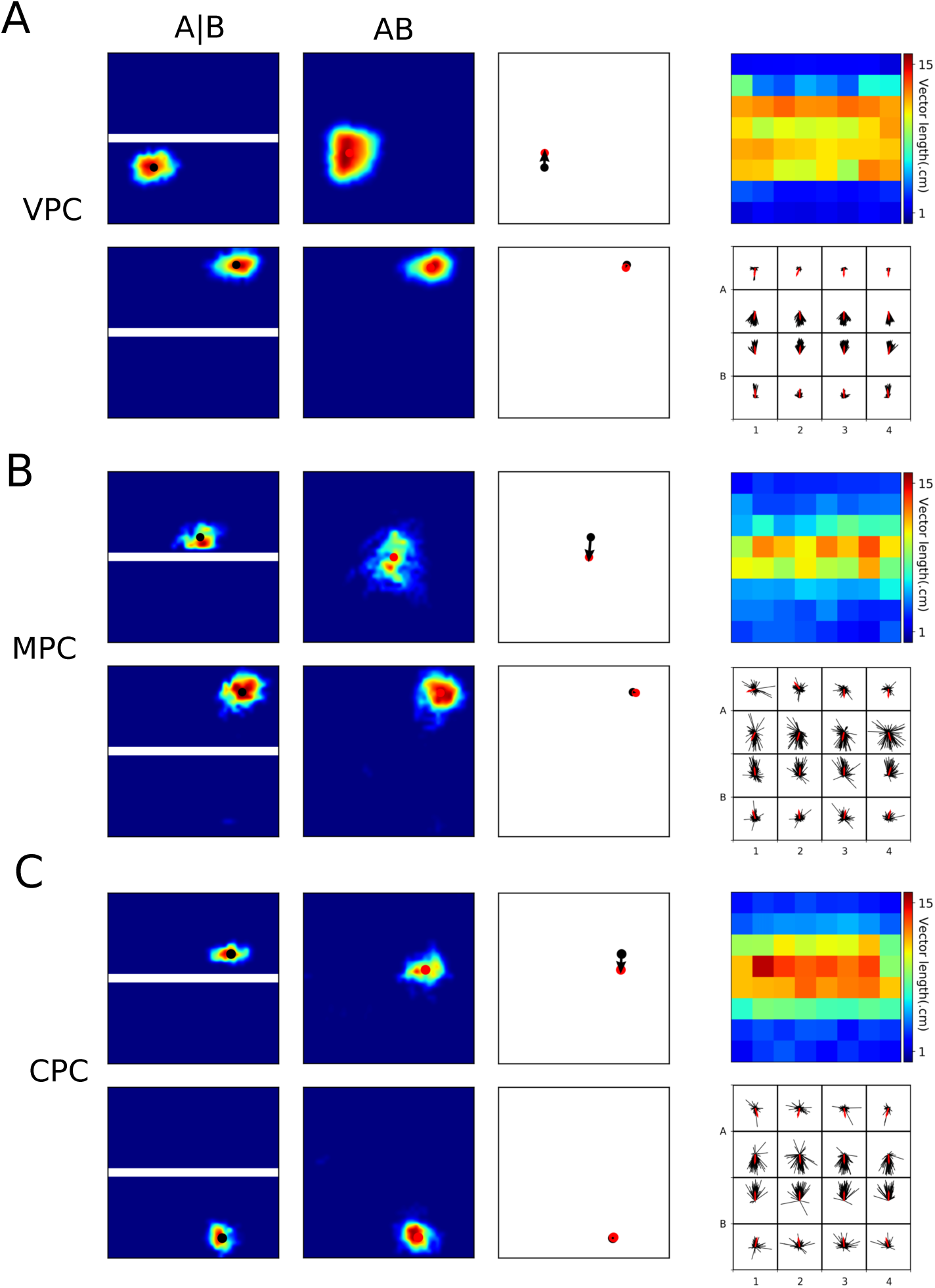
Place fields in the merged-room experiment. A. Left: receptive fields of two VPCs in the training and testing environments, either close to the removed wall (top) or distal from it (bottom). Middle: displacement vectors of the cells on the left. Right: color map of displacement vector lengths for all cells (top) and all displacement vectors with their mean direction shown in red (right). B,C. Receptive fields and displacement vectors for MPCs (B) and CPCs (C). Refer to A for details.

Thus, the low-correlation band near the location of the removed wall is induced in the model by changes in visual input in the merged environment, which affect place coding via VPC activities. Local visual features at the locations distant from the removed wall are similar in the corresponding locations of the original environments A and B, since visual patterns formed by the closest walls and extramaze cues remain largely unchanged after the central wall removal. Therefore, VPCs activities at these locations during testing are very similar to those during training (Fig. 5A), leading to the same grid pattern at these locations. However, at the locations close to the removed wall, the combined effect of stable distal cues and modified proximal wall cues result in an extension of VPC receptive fields over the previous location of the removed wall. These changes in visual receptive fields induce local corrections of grid cell activity by shifting grid-cell activity packets towards the center, resulting in local deformations of grid-cell firing patterns similar to those observed during gain modification experiments. These deformations will in turn affect place fields of MPCs and CPCs, by shifting place fields of the cells near the removed wall towards it (Figs. 5B,C). These results suggest that local deformations of grid fields can result from the same correction mechanism as the one studied in the previous section, but in which local sensory conflict is induced by changes in the visual input instead of changes in self-motion gain.

Two principal neural processes affect the formation of spatial representation in our model: while the acquisition of new spatial representations crucially depends on synaptic plasticity, the dynamic interaction between visual and self-motion cues is mediated by neuronal dynamics. We therefore tested the contribution of these two processes to the observed results. The influence of plasticity was assessed by letting the model learn during testing in the merged room, while that of neuronal dynamics was tested by progressively decreasing the strength of the loop (i.e. decreasing the control of vision over self-motion cues) in the absence of synaptic plasticity. The results of these manipulations can be summarized as follows. First, when learning was allowed during testing and the testing session in the merged room was sufficiently long, the particular correlation pattern (see Fig. 4C,D) was broken and a new representation was formed as a result of learning (Figs. 6A-C), unlike what was observed by Wernle et al. In particular, the newly formed global pattern was aligned with only one of the walls, resembling the results of Carpenter et al. (2015) addressed in the following section. Moreover, learning of the new representation was faster when the control of visual cues (controlled by *α*) was low (not shown), since slower dynamics favors the learning of new connections between self-motion-based and visual representations. These results suggests that either the band-correlation pattern is a transient effect and should disappear with a longer exposure to the environment, or that learning of a new representation is inhibited in the merged room in real rats. Second, the decrease of *α* across separate sessions resulted in widening of the low correlation band (Fig. 6D). This modification of the correlation pattern is explained by the fact that under a weak control of place fields by vision, it takes longer for the visual cues to correct self-motion.

**Figure 6:**
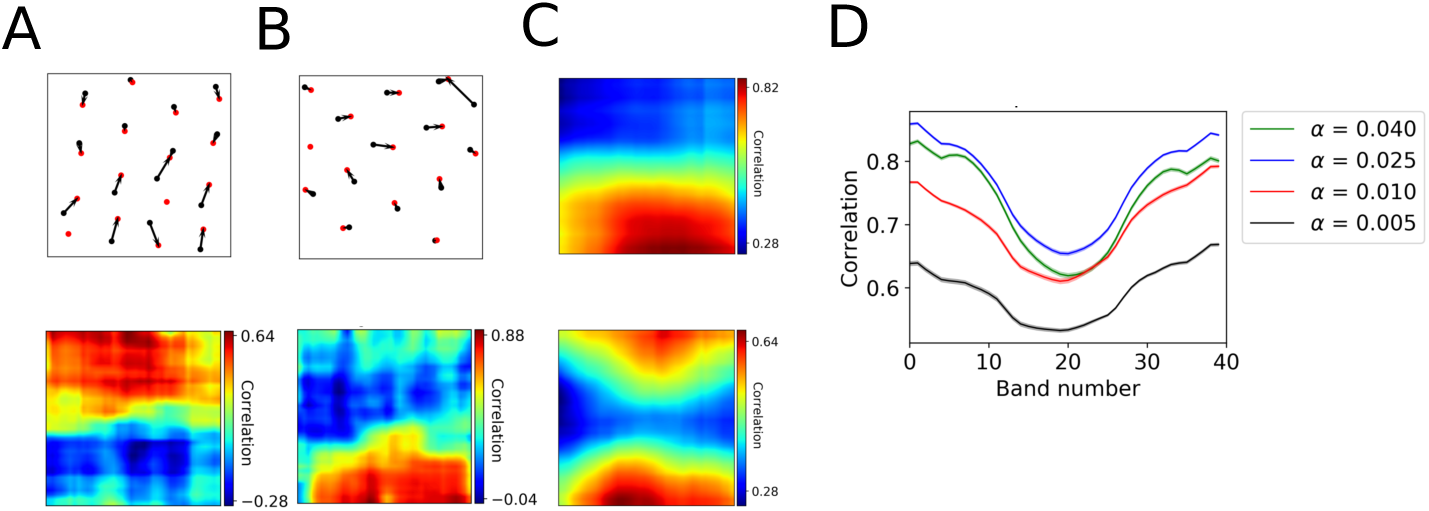
Influence of plasticity and dynamics on grid patterns in the merged-room experiment. A,B. Displacement vectors (top) and corresponding sliding correlation maps (bottom) of two example grid cells after learning in the merged room. C. Averaged over many grid cells, sliding correlation maps can result in different mean correlation patterns. D. Correlation profile for different values of the the strength *α* of the hippocampal feedback loop.

### 3.2. Double-room experiment

In the experiment of Carpenter et al. (2015), grid cells were recorded in rats during foraging in an experimental environment consisting of two rectangular rooms connected by a corridor (Fig. 7A, see Carpenter et al., 2015). The rooms were rendered as similar as possible in their visual appearance in order to favor visual aliasing. If local visual cues are the main determinant of grid cell activity, identical grid fields in the two environments were expected. In contrast, if self-motion cues are used to distinguish between the two rooms, grid cells should have distinct firing fields in the two environments. The results of this experiment revealed that both external and internal cues influence neuronal activity, but in a temporally-organized fashion. In particular, during early exploration sessions, grid cells had similar firing patterns in the two rooms, and this effect was maintained during the whole period of a session (tens of minutes). However, as the number of sessions (or days, as 1 session per day was run) increased, grid cells formed a global representation of the experimental environment, such that initial association between local cues and grid fields was progressively lost in one of the two rooms. These results are in apparent conflict with the data from the merged-room experiment considered earlier, since in that experiment local cues at the distant walls kept their control of nearby grid fields for up to 10 daily sessions.

**Figure 7:**
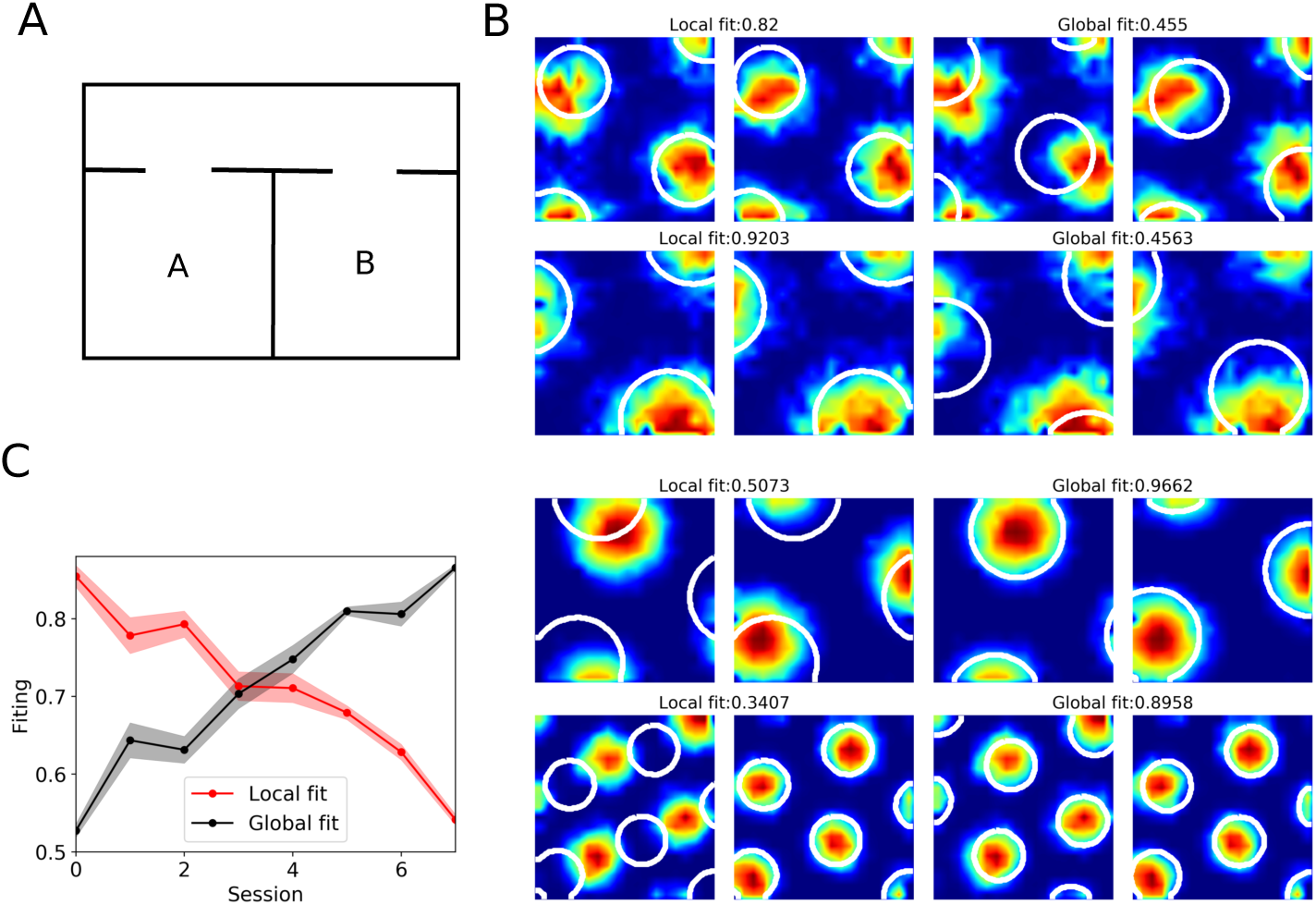
Simulation of the double-room experiment of Carpenter et al., 2015. A. Top view of the experimental environment. B. Local fit (left) versus global fit (right) during early (top) and late (bottom) sessions for two example grid cells (rows). C. Population estimates of the local fit (red) and global fit (black) as a function of session number (the value of *α* decreased from 0.04 to 0.005 across sessions).

What could be the reason for the differences in learned grid-cell representations in the two experiments? Suppose that, in the conditions of the double-room paradigm, the rat first enters room A, such that initial associations between self-motion and visual cues are established in that room. The key question is whether or not a new representation for the subsequently entered room B will be formed, despite its identical visual appearance with room A (note that in the following we refer to any initially experienced room as room A, independently on which actual room was visited first in the simulations). Results from the previous section suggest that a weaker control of visual cues combined with synaptic plasticity leads to the formation of a new representation. To verify this hypothesis, we run our model in the conditions of Carpenter et al. experiment, and we progressively (i.e. session by session) decreased the strength of the hippocampal-entorhinal feedback loop (without disabling synaptic plasticity). As the feedback strength controls the influence of visual input in our model, we expected that this procedure will result in the construction of a global representation on the level of grid cells when the strength of the loop is sufficiently low. This was indeed the case as the global fit was high when the loop strength was set to low values (small *α*), and, conversely, the local fit was high for a strong loop (Figs. 7B,C, both of these measures were calculated in the same way as in the study by Carpenter et al., 2015, see also Appendix).

The local representation in early sessions is a consequence of the fact that only a representation of one room is learned, so that once the model rat enters the second room, grid-cells activities are quickly reset by vision to the representation of the first (or, in terms of Skaggs and McNaughton (1998), the representation of room A is “instantiated” upon the entry to the room B). In this case both MPCs and CPCs had identical firing fields in the two rooms (Fig. 8A). This was quantified in the model by computing the spatial correlation between place fields of each cell in the two rooms (correlation of 1 corresponds to identical place fields). On the level of the whole population, the mean place-field correlation is high for a strong feedback loop (early sessions, large *α*, Fig. 8B). The transition to a global representation in later sessions results from newly formed synaptic associations between MPCs in CA3 (that are under a strong influence of self-motion input from grid cells), and CPCs in CA1 that are driven by vision. Synaptic plasticity at these connections is favored by a decreased hippocampal input to the EC, leading to a stronger reliance on self motion (late sessions, small *α*, Fig. 8B). The development of such a new representation is reflected in lower place-field correlation on the level of MPCs and CPCs (late sessions, small *α*, Fig. 8B). Note that purely vision-driven VPCs always have identical place fields in the two environments (not shown).

**Figure 8:**
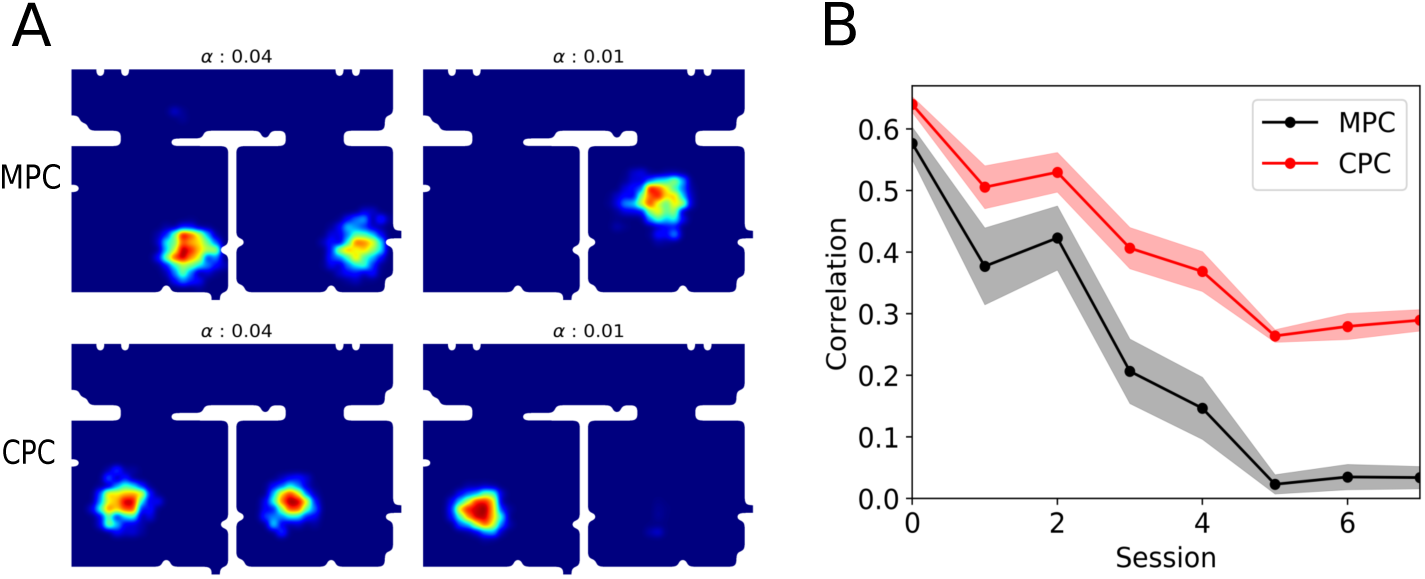
Evolution of place fields in the double room experiment. A. An example of MPC (top) and CPCs (bottom) place field during early learning sessions (left column, high *α*) and late sessions (right column, low *α*). In early sessions a majority of place cells have similar place fields in the two rooms, whereas in late sessions a majority of place cells have a place field only in one of the rooms. B. Spatial correlation between place fields of a cell in the two rooms, averaged over all place cells, as a function of session number (or, equivalently, as a function of decreasing value of *α*.

To summarize, the results of both the merged-room experiment of Wernle et al. (2018) and the double-room experiment of Carpenter et al. (2015) can be explained by the same model under two assumptions: First, synaptic plasticity is slow or inhibited when rats are placed into the merged room after learning in room A and B, but not when the rats are exposed to a stable double-room environment; Second, the control of visual cues progressively decreases in a familiar environment in the course of daily sessions (this requirement is crucial to reproduce the result of the second experiment, but, according to our simulations, has only a weak effect in the first). What could be the explanation for the inhibition of learning in the merged-room, as opposed to continuous learning in the double-room experiment across daily sessions? Analysis of our model offers the following possible explanation: In early sessions of the double-room experiment, a large mismatch between visual (i.e. encoded in VPC activities) and self-motion (encoded by MPC activities) input occurs at the moment of entry to, or exit from, the room B, since the population activity of VPSs “jumps” to reflect the room A cues or the corridor cues, respectively. This jump of population activity can be quantified by the drop in correlation between the projections of VPCs and MPCs in CA3 onto the CPCs in CA1 near the room doors (Fig. 9A). In contrast, the mismatch is smaller for the merged-room experiment, since the visual and self-motion cues near the removed wall code for similar spatial positions (Fig. 9B). Therefore, it is possible that learning across sessions is regulated by the size of the mismatch between visual and self-motion cues. Note that statistical characterization of the mismatch in Fig. 9 required averaging over many experimental runs and even in our idealized model can not be reliably detected online. This could be a possible reason why building of a global environment representation in Carpenter et al. experiment takes many days. We thus propose that CA1 area or, more likely, its output structures implement a mismatch detection process that can regulate hippocampal synaptic plasticity on the time scale of days (see below).

**Figure 9:**
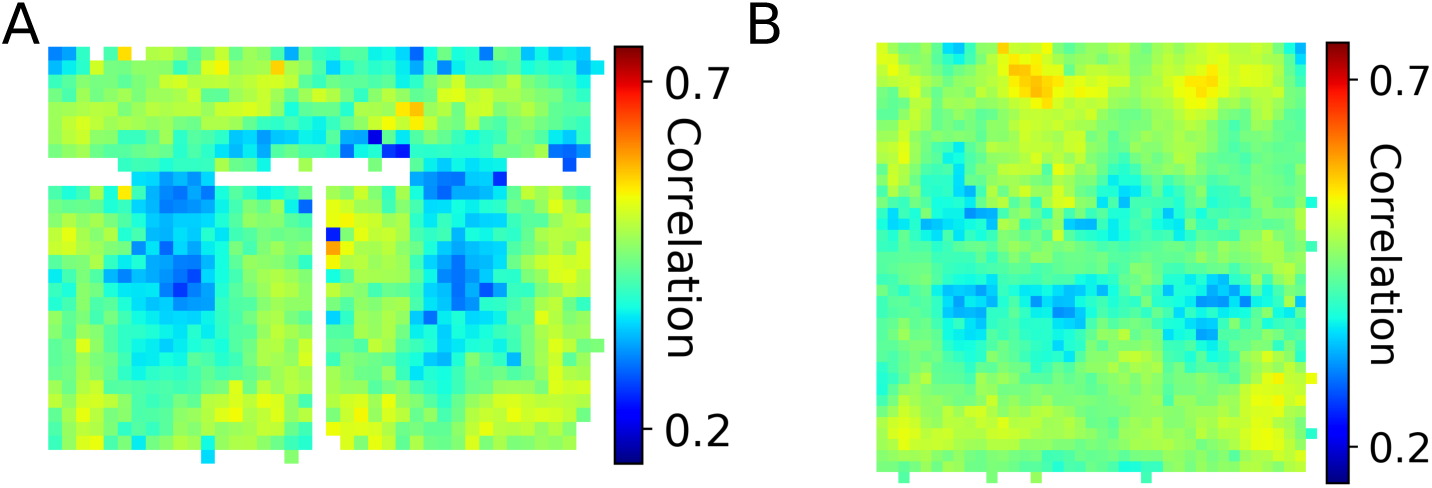
Mismatch between the visual and self-motion representations in the double-room (A) and merged-room (B) experiments. The colors denote the correlation between VPCs and MPCs projections onto the CPS population.

## 4. Discussion

Our model is based on two main assumptions: that of a loop-like dynamics in the entorhinal-hippocampal network, and that of an independent visual place-cell representation formed on the basis of hippocampal inputs other than grid cells. Place cells in the CA1 area receive spatially organized inputs from grid cells via direct projections from mEC layer III and via perforant path projections from layer III via DG and CA3. Isolating direct feedforward mEC input to CA1 only weakly affect place-sensitive activity in CA1 suggesting that mEC inputs are sufficient for the establishment of representation in this area (Brun et al., 2002). Isolating only indirect projections resulted in noisier CA1 place fields that formed although relatively impaired but still stable spatial representations (Brun et al., 2008). These results suggest a complementary role of both the direct and indirect pathways for spatial coding in the CA1. Place cells in CA1 project back to the entorhinal cortex both directly and via subiculum (Naber et al., 2001, Kloosterman et al., 2003, Slomianka et al., 2011) and hippocampal input is necessary for grid cell activity (Bonnevie et al., 2013), supporting the loop-like structure of entorhinal-hippocampal interactions (Iijima et al., 1996). That a subset of hippocampal place cells can form spatial representations independently from grid cells is supported by the evidence showing that place fields can exist before the emergence of the grid cell network in rat pups (Muessig et al., 2015) and that the disruption of grid cell activity in adult rats does not prevent the emergence of place fields in novel environments (Brandon et al., 2014). These grid-cell independent place fields retain all principal properties of a self-organized representation in control animals: it can be learned in new environments, it is stable over time, and independent maps are established in different rooms (Rueckemann et al., 2016, Schlesiger et al., 2018). These data suggest that some place cells rely mostly on grid-cell input, likely representing self-motion-based spatial signals, while other place cells preferentially use other sensory information to form spatial representation in a self-organized manner. This separation of place cells depending on their principal source of sensory input is also supported by observations showing that in virtual environment subsets of place cells are differentially responsive to sensory manipulations: during passive movement 25% of cells keep their firing fields unchanged; 20% of cells do change their firing patterns when all visual cues are turned off; most of the cells are modified to various degrees by cue manipulations (Chen et al., 2013), see also Markus et al. (1994). Moreover, recent evidence suggests that CA1 cells responsive to visual and self-motion input are anatomically separated: place cells more responsive to self-motion cues are located predominantly in superficial layers of CA1, while those more responsive to visual cues are found in deep layers (Fattahi et al., 2018), see also Mizuseki et al. (2011). It was also recently shown that CA1 cells in deep and superficial layers receive stronger excitation from mEC and lEC, respectively, with the amount of excitation being also dependent on the position of the neurons along the longitudinal hippocampal axis (Masurkar et al., 2017). These data further support the existence of functionally different subsets of place cells in CA1, that can either be inherited from similarly segregated cells in CA3 or to be formed directly from lEC inputs to CA1.

Our model is constructed to reflect the above data in a simplified way. While the neural basis for the aforementioned grid-cell-independent code is not clear, we conceptualized it by a population of VPCs, which learn subsets of visual features corresponding to a particular location using simple competitive learning scheme. Similarly to experimental data described above, VPCs form a stable and independent code for different environments as long as visual cues in these environments are stable. It is likely that such a code is formed inside the hippocampus itself based on the inputs either from parietal-cingulate network (Byrne et al., 2007, Bicanski and Burgess, 2018), or from lEC input (Schlesiger et al., 2018), since no location-sensitive code was observed directly upstream of the hippocampus (but see Mao et al., 2017). While in its current version our model assumes that VPCs are learned in CA3 and transmitted to CA1, the model can be modified to implement competitive learning in CA1 directly on visual inputs from lEC, bypassing CA3. Our self-motion based code in GC-MPC populations is based on internal attractor dynamics and does not in principle require place-cell input, contrary to experimental data (Bonnevie et al., 2013). However, this dependence can be included in the model by adding strong inhibition to the grid cell layer, such that a nonspecific excitatory drive from CA1 were required for grid-cell activities (Bonnevie et al., 2013). Such a modification of the model will not significantly change any of the present results.

Main conclusions from our modeling results are twofold. First, the construction of a global representation in the double-room experiment requires a diminished control of visual cues over path integration, translated in the model by decreasing the strength of the hippocampal input to the EC. By slowing down the dynamical correction of GCs and MPCs by vision, it allows synaptic plasticity to form new associations between visual representations (encoded in VPC activity) and CA3-mediated representations at the level of CA1, and to disambiguate the two rooms. Thus, in our model, synaptic plasticity at CA3-CA1 synapses is crucial for the formation of new representations in visually identical environments. Ultimately, the construction of this representation is determined by relative time scales of two processes: *(i)* correction of path integration by visual cues using network dynamics, and *(ii)* synaptic plasticity at Schaffer collaterals. Second, the fact that rats learn a global representation in the double-room, but not in the merged-room experiment is explained in the model by a strongly reduced or inhibited synaptic plasticity in the latter case. Indeed, under the hypothesis that grid cells express hexagonal patterns as a consequence of attractor dynamics with circular weight matrices (McNaughton et al., 2006), translocation of grid fields at the center of the environment must result from dynamic correction mechanisms, since synaptic plasticity between place-cell and grid-cell networks will necessarily lead to the emergence of a coherent (global) grid-cell representation. If this explanation is correct, then what could be the mechanism that regulate synaptic plasticity differently in the two cases? One possibility suggested by the analysis of the model is that such a regulation mechanism can act on the basis of a mismatch between visual and self-motion representations. On the level of population activity, a high degree of mismatch corresponds to incoherent “jumps” of visual representation caused by visual aliasing, relative to the representation formed by path integration. While these jumps are reflected in the distribution of synaptic inputs to modeled CA1 cells in our model (Fig. 9), the fact that learning of a global representation in real animals takes many days (Carpenter et al., 2015) suggests that detection of this mismatch may involve memory consolidation mechanisms (Skaggs and McNaughton, 1996, Girardeau et al., 2009, Benchenane et al., 2010).

A number of experiments studied place fields dynamics in environments consisting of two or more visually identical compartments (Skaggs and McNaughton, 1998, Tanila, 1999, Fuhs et al., 2005, Paz-Villagŕan et al., 2006, Spiers et al., 2015, Grieves et al., 2016). The objective of these experiments was to check whether path integration can be used to distinguish between compartments and to assess the extent to which visual cues control path integration information. Earlier experiments provided evidence for a partial (Skaggs and McNaughton, 1998) or a nearly complete (Tanila, 1999) remapping when rats travelled between two similarly looking compartments, suggesting that path integration can be used to distinguish between them. A major difference between experimental setups in these latter experiments was that the two compartments in Skaggs and McNaughton (1998) were oriented in the same way, whereas in Tanila (1999) there was a 180° difference in their orientation. A follow-up experiment (Fuhs et al., 2005) has demonstrated a key role of angular, but not linear, path integration in complete remapping observed by Tanila et al. 1999. However, Fuhs et al. did not observe partial remapping in conditions very similar to those of Skaggs and McNaughton (1998), as most cells had identical place fields in the two compartments. More recent experiments with multiple visually identical compartments confirmed the importance of angular path integration for remapping (Spiers et al., 2015, Grieves et al., 2016, see also Paz-Villagŕan et al., 2006), and suggested that a long amount of time (about 2-3 weeks) is necessary to build separate representations for visually identical rooms connected by a corridor (Carpenter et al., 2015).

In our simulations, we assumed that the animals head direction system provides a correct orientation information (i.e. relative to an arbitrary fixed reference orientation) at any moment in time, and so the visual input to the model is always aligned to the common directional frame in all environments (in the experiment of Carpenter et al. a common directional frame could be provided by the corridor cues, whereas it was provided by distal extramaze cues in Wernle et al. experiment). As a result of competitive learning, synapses to a visual place cell learn visual cues observed at a location where this cell was recruited. Therefore, a place is visually “recognized” (i.e. visual place cells strongly fire) if the previously learned visual cues are observed in the same allocentric direction (independently of any path integration signal). If, however, the same visual cues are observed at a very different orientation (e.g. is a room is rotated 180°) visual place cells will not be activated (unless visual cues are rotationally symmetric), and new cells will be recruited to represent this environment, in agreement with Fuhs et al. (2015) study. At smaller rotation angles, the model predicts that place cells will be activated to a higher degree, depending on the autocorrelation width of the learned visual snapshots (Grieves et al., 2016). That the head direction system can maintain a fixed orientation in the presence of visual cue rotation is supported by experimental evidence (Jacob et al., 2017, see also Paz-Villagŕan et al., 2006).

The ability (or inability) of the hippocampal representations to express partial remapping has been discussed in view of the multichart model (McNaughton et al., 1996, Samsonovich and McNaughton, 1997). This model predicted that if rats could learn room identities despite their similar visual appearance, place-field representations of the two rooms would be orthogonal (different charts are active in different rooms), whereas they would be identical in the opposite case (the same chart is active in both rooms). Partial remapping observed by Skaggs and McNaughton (1998) contradicted this hypothesis, as some cells had identical fields in the two rooms, while other cells and different place fields, suggesting that two charts could be active at the same time. In similar conditions Fuhs et al. (2005) observed no partial remapping for unclear reasons, but suggested that the map of one compartment was somehow “extended” to the second one, instead of loading a new map. Our results contribute to this question in two ways. First, we argued that a learning of new representation is under control of a putative neural mismatch detection mechanism. In the experimental conditions of the two above studies, the largest amount of mismatch occurs upon the door crossing, and so the number of door crossings experienced by the rat may be an important parameter with respect to learning. While in Skaggs and McNaughton (1998) the rats were freely moving between the compartments during a trial, in Fuhs et al. (2005) the number of transitions between rooms was limited to 2 per trial, potentially affecting the results. Second, our results provide a neuronal mechanism for the map observed map extension, i.e. progressive learning of a global representation.

Our results lead to a number of testable predictions. First, VPC in the model acquire representation of only one compartment (among two or more identically looking ones). We thus predict that a subset of place cells, that do not rely on self-motion signals (e.g. such as those observed in Chen et al., 2013) and potentially located in the deep sublayer of CA1 pyramidal layer (Fattahi et al., 2018), will persist through learning and will have repetitive place fields even when a global representation has been learned. Second, learning of separate neuronal representations of different compartments (i.e. progressive remapping) will be require the formation of new associations between CA3 cells and CA1 cells preferentially from the superficial sublayer of pyramidal cells. Third, place cells that will remap first should have place fields close to the door, since for these cells the difference between visual and motion-based inputs is largest. Finally, as the width of the low-correlation band (Fig. 6D) is proposed to be related to the strength of the visual cue control over path integration, it is predicted that stronger reliance on path integration will result in a wider band. This might occur for example in aged animals, in which a stronger reliance on path integration (or, conversely, an weaker control by visual cues) has been observed (Tanila, 1999, Rosenzweig et al., 2003).

## Acknowledgements

Funding: This research was supported by ANR – Essilor SilverSight Chair ANR-14-CHIN-0001.

## Appendix

### Visual input

The artificial retina was modeled as a rectangular grid of Gabor filters uniformly covering the panoramic cylindrical camera with visual field 160° x 360°. At each location of the grid, 4 filters of different orientations were used. We used two spatial frequencies for all the filters (180 Hz, 72 Hz) chosen so as to detect visual features of simulated environments. Activities of all Gabor filters were computed by the convolution with the input visual image at each time step. Filter activities were then aligned with a common allocentric directional frame, such that if the model rat rotated without changing its spatial position, the activities of aligned filters would stay constant.

### Virtual environments

Virtual environments for the three simulations presented in this paper were developed with Unity (www.unity3d.com). In Simulation 1 (Figs. 2 and 3) the environment was a rectangular room 2×1 m with featureless gray walls. In Simulation 2 (Figs. 4-6), the experimental room was modeled as a square arena 2×2 m. During training, it was separated into two rooms by a wall at the center of the environment. The experimental arena was located inside a bigger environment (4×4 m) with four salient visual cues (large circles) on each wall. In Simulation 3 (Figs. 7-8), the environment consisted of two identical rooms 1×1 m connected by a corridor (0.5×2 m).

#### Simulation details

In all three simulations, VPCs were learned from the simulation environment before the training of the place cells and grid cells. Model parameters are listed in Table 1.

**Table 1:**
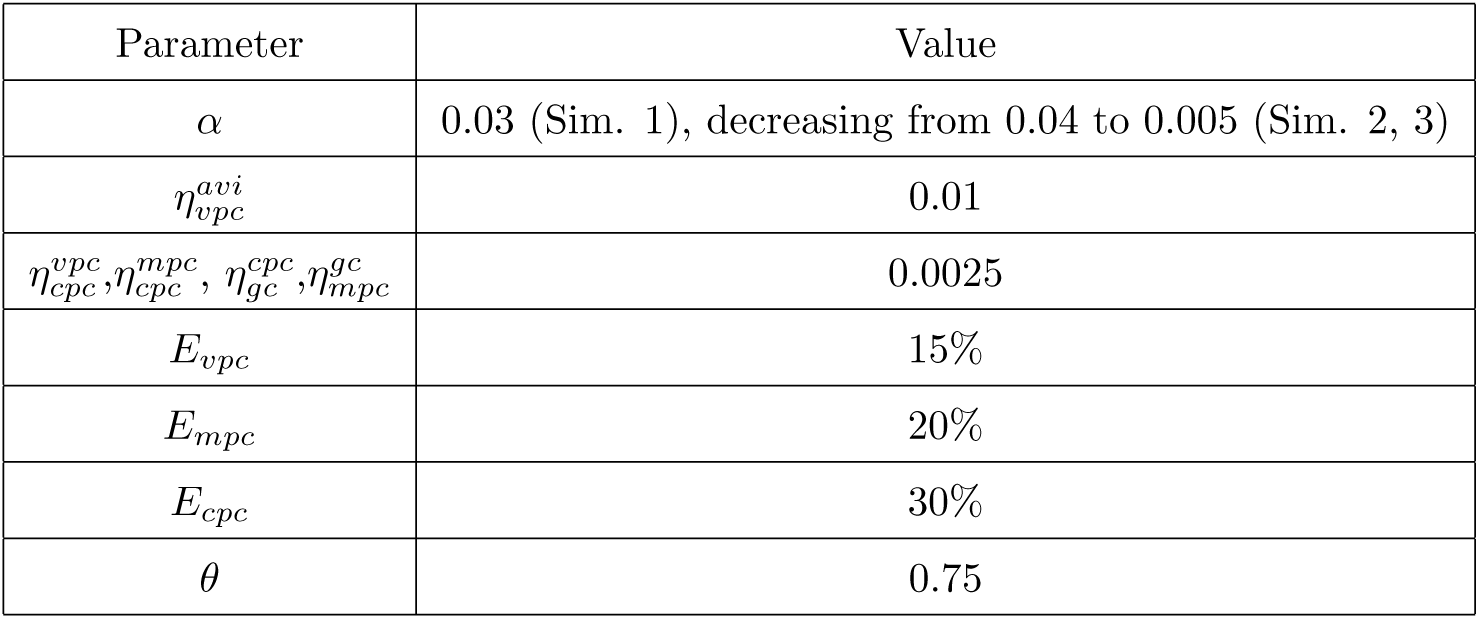
Parameters of the model.

### Simulation 1

#### Training

The model was trained for about 25 minutes (15000 time steps) by moving quasi-randomly in the experimental room.

#### Testing

Synaptic weights were fixed, and activities of all the cells in the model were recorded in the following three experimental conditions. In the ‘light’ condition the full model was run to randomly explore the environment. In the ‘passive translation’ condition, the velocity vector input to the grid cell populations was set to (0, 0). In the ‘dark’ condition, the model was run with visual cues turned off (uniform gray images were presented as visual input). Next, the trained model was run to cross the environment from left to right in ‘light’ and ‘dark’ conditions as before, but with the speed gain in the grid cell populations modulated as described in the Results.

### Simulation 2

#### Training

The model was trained separately in rooms A and B for 30 minutes, and synaptic weight were fixed to the learned values.

#### Testing

In the main experiment, neural activities were recorded while the model rat randomly explored the merged room for 1 h. In the experiment testing the influence of plasticity, synaptic weights were updated while the model rat additionally explored the merged room for 1h. In the experiment testing the influence of the strength of the feedback loop, the model rat was run in the merged room for 20 trials per each value of *α*, ranging from 0.005 to 0.04. To average data, 4 testing trials were run in each condition.

#### Sliding correlation

The sliding correlation heat maps for grid-cell firing patterns were calculated as described in Wernle et al. (2018). The size of the sliding correlation window was defined based on the grid spacing of the cell. The window moved from the top left to the bottom right corner in the grid field maps of the environment A|B (i.e. before the wall removal) and AB (i.e. after the wall removal). At each window location, the portion of the grid maps in the environments A|B and AB, outlined by the sliding window, were correlated with each other.

#### Displacement vector analysis

Displacement vectors were calculated as described in Wernle et al. (2018). To obtain a displacement vector for one grid cell, the experimental environment was divided into 4×4 blocks (50×50 cm each). In each block, the vector corresponding to the shift of grid fields in the environment AB relative to that in the environment A|B was calculated. The vectors were sorted into the corresponding blocks based on the grid field location in the training environment and the mean over all vectors was computed. To analyze displacement vector lengths, the environment was divided into 8×8 bins. The vectors were then sorted into the corresponding bins based on the original grid field location in the training environment, and the mean vector length was computed.

### Simulation 3

#### Training

At the beginning of each training session, the model was placed into the center of the corridor and then explored the complete environment quasi-randomly for 1 h. In subsequent training sessions, the strength of the feedback loop *α* decreased from 0.04 (first session) to 0.005 (last session) with step 0.005.

#### Testing

After each training session, the weights were fixed and neural activity was recorded. In order to average the results, the experiment was repeated 20 times for each value of *α*.

#### Global and local fits

The firing rate maps of modeled grid cells were fit with ideal local and global grid patterns using the procedure described in Carpenter et al. (2015). First, grid spacing was identified by correlating the firing pattern with 30 ideal firing grids. Each ideal grid pattern is a product of three cosine gratings

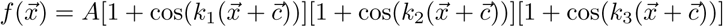

with peak firing rate *A*, wave vectors 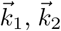 and 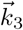 and phase offsets 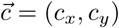. The wave vectors are defined as 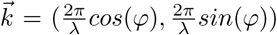, where 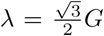 is the grating wave length, *G* is the grid spacing and *ϕ* is the grid orientation. The 30 ideal grid patterns were created with grid spacing evenly distributed between 30 and 170 cm. Since the grid orientation in the model is set to 7.5°, *ϕ* in the three wave vectors is equal to 7.5°, 127.5° and 247.5°, respectively. Spatial cross-correlograms were computed between the recorded firing rate map and the ideal grid patterns over a range of spatial phase offsets. The grid spacing of the recorded firing pattern is then set to that of the ideal grid pattern with the highest correlation. Second, a local and global fit with the identified grid spacing was computed for the recorded firing rate map. The local fit was performed using two grid patterns (one per room) with the same phase offset. The global fit was performed using only one grid pattern with continuous phase across the two rooms. The Pearson product-moment correlation between the recorded firing rate map and the local and global grid patterns were computed over a range of phase offsets. The highest correlation with the local and global model was identified as the value of local and global fit, respectively.

## References

J. O’Keefe, L. Nadel, The hippocampus as a cognitive map, Clarendon Press, Oxford, ISBN 0198572069, 1978.

J. O’Keefe, J. Dostrovsky, The hippocampus as a spatial map. Preliminary evidence from unit activity in the freely-moving rat, Brain Res. 34 (1971) 171–175.

J. O’Keefe, A. Speakman, Single unit activity in the rat hippocampus during a spatial memory task, Exp. Brain Res. 68 (1987) 1–27.

R. U. Muller, J. L. Kubie, The effects of changes in the environment on the spatial firing of hippocampal complex-spike cells, J. Neurosci. 7 (7) (1987) 1951–1968, ISSN 0270-6474.

J. J. Knierim, H. S. Kudrimoti, B. L. McNaughton, Interactions between idiothetic cues and external landmarks in the control of place cells and head direction cells., J. Neurophysiol. 80 (1) (1998) 425–46.

R. P. Jayakumar, M. S. Madhav, F. Savelli, H. T. Blair, N. J. Cowan, J. J. Knierim, Recalibration of path integration in hippocampal place cells, bioRxiv (2018) 319269 doi:10.1101/319269.

E. J. Markus, C. A. Barnes, B. L. McNaughton, V. L. Gladden, W. E. Skaggs, Spatial information content and reliability of hippocampal CA1 neurons: Effects of visual input, Hippocampus 4 (4) (1994) 410–421, ISSN 1050-9631, doi:10.1002/hipo.450040404.

G. Chen, J. A. King, N. Burgess, J. O’Keefe, How vision and movement combine in the hippocampal place code., Proc. Natl. Acad. Sci. U. S. A. 110 (1) (2013) 378–383, ISSN 1091-6490, doi:10.1073/pnas.1215834110.

M. Fattahi, F. Sharif, T. Geiller, S. Royer, Differential Representation of Landmark and Self-Motion Information along the CA1 Radial Axis: Self-Motion Generated Place Fields Shift toward Land-marks during Septal Inactivation, J. Neurosci. 38 (30) (2018) 6766–6778, ISSN 0270-6474, doi:10.1523/JNEUROSCI.3211-17.2018, URL http://www.jneurosci.org/lookup/doi/10.1523/JNEUROSCI.3211-17.2018.

M. Fyhn, S. Molden, M. P. Witter, E. I. Moser, M. B. Moser, Spatial representation in the entorhinal cortex., Science (80-.). 305 (2004) 1258–1264.

B. L. McNaughton, F. P. Battaglia, O. Jensen, E. I. Moser, M. B. Moser, Path integration and the neural basis of the ‘cognitive map’, Nat. Rev. Neurosci. 7 (8) (2006) 663–678.

R. M. Hayman, K. J. Jeffery, How heterogeneous place cell responding arises from homogeneous grids-A contextual gating hypothesis, Hippocampus 18 (12) (2008) 1301–1313, ISSN 10509631, doi:10.1002/hipo.20513.

S. Cheng, L. Frank, The structure of networks that produce the transformation from grid cells to place cells, Neuroscience 197 (2011) 293–306, ISSN 0306-4522, doi:10.1016/J.NEUROSCIENCE.2011.09.002.

T. Hafting, M. Fyhn, S. Molden, M. B. Moser, E. I. Moser, Microstructure of a spatial map in the entorhinal cortex., Nature 436 (2005) 801–806.

J. Krupic, M. Bauza, S. Burton, C. Barry, J. O’Keefe, Grid cell symmetry is shaped by environmental geometry, Nature 518 (7538) (2015) 232–235, ISSN 0028-0836, doi:10.1038/nature14153.

M. Fyhn, T. Hafting, A. Treves, M.-B. Moser, E. I. Moser, Hippocampal remapping and grid realignment in entorhinal cortex., Nature 446 (7132) (2007) 190–4, ISSN 1476-4687, doi:10.1038/nature05601.

T. Sasaki, S. Leutgeb, J. K. Leutgeb, Spatial and memory circuits in the medial entorhinal cortex, Curr. Opin. Neurobiol. 32 (2015) 16–23, ISSN 18736882, doi:10.1016/j.conb.2014.10.008.

M. I. Schlesiger, B. L. Boublil, J. B. Hales, J. K. Leutgeb, S. Leutgeb, Hippocampal Global Remapping Can Occur without Input from the Medial Entorhinal Cortex, Cell Rep. 22 (12) (2018) 3152–3159, ISSN 22111247, doi:10.1016/j.celrep.2018.02.082, URL https://linkinghub.elsevier.com/retrieve/pii/S2211124718302924.

T. Wernle, T. Waaga, M. Mørreaunet, A. Treves, M. B. Moser, E. I. Moser, Integration of grid maps in merged environments, Nat. Neurosci. 21 (1) (2018) 92–105, ISSN 15461726, doi:10.1038/s41593-017-0036-6.

F. Carpenter, D. Manson, K. Jeffery, N. Burgess, C. Barry, Grid Cells Form a Global Representation of Connected Environments, Curr. Biol. 25 (9) (2015) 1176–1182, ISSN 09609822, doi:10.1016/j.cub.2015.02.037.

T. Solstad, E. I. Moser, G. T. Einevoll, From grid cells to place cells: A mathematical model, Hippocampus 16 (12) (2006) 1026–1031.

J. O’Keefe, N. Burgess, Dual phase and rate coding in hippocampal place cells: Theoretical significance and relationship to entorhinal grid cells, Hippocampus 15 (7) (2005) 853–866, ISSN 10509631, doi:10.1002/hipo.20115.

H. T. Blair, K. Gupta, K. Zhang, Conversion of a phase- to a rate-coded position signal by a three-stage model of theta cells, grid cells, and place cells, Hippocampus 18 (12) (2008) 1239–1255, ISSN 10509631, doi:10.1002/hipo.20509.

D. Sheynikhovich, R. Chavarriaga, T. Strösslin, A. Arleo, W. Gerstner, T. Strosslin, A. Arleo, W. Gerstner, Is there a geometric module for spatial orientation? Insights from a rodent navigation model., Psychol. Rev. 116 (3) (2009) 540–566, ISSN 0033295X, doi:10.1037/a0016170.

P. K. Pilly, S. Grossberg, How Do Spatial Learning and Memory Occur in the Brain? Coordinated Learning of Entorhinal Grid Cells and Hippocampal Place Cells, J. Cogn. Neurosci. 24 (5) (2012) 1031–1054, ISSN 0898-929X.

T. Bonnevie, B. Dunn, M. Fyhn, T. Hafting, D. Derdikman, J. L. Kubie, Y. Roudi, E. I. Moser, M.-B. Moser, Grid cells require excitatory drive from the hippocampus, Nat. Neurosci. 16 (3) (2013) 309–317, ISSN 1097-6256, doi:10.1038/nn.3311, URL http://www.nature.com/articles/nn.3311.

V. H. Brun, S. Leutgeb, H.-Q. Wu, R. Schwarcz, M. P. Witter, E. I. Moser, M.-B. Moser, Impaired Spatial Representation in CA1 after Lesion of Direct Input from Entorhinal Cortex, Neuron 57 (2) (2008) 290–302, ISSN 0896-6273, doi:10.1016/j.neuron.2007.11.034.

J. W. Rueckemann, A. J. DiMauro, L. M. Rangel, X. Han, E. S. Boyden, H. Eichenbaum, Transient optogenetic inactivation of the medial entorhinal cortex biases the active population of hippocampal neurons, Hippocampus 26 (2) (2016) 246–260, ISSN 10509631, doi:10.1002/hipo.22519, URL http://doi.wiley.com/10.1002/hipo.22519.

L. Muessig, J. Hauser, T. J. Wills, F. Cacucci, A Developmental Switch in Place Cell Accuracy Coincides with Grid Cell Maturation, Neuron 86 (5) (2015) 1167–1173, ISSN 0896-6273, doi:10.1016/J.NEURON.2015.05.011.

J. Koenig, A. N. Linder, J. K. Leutgeb, S. Leutgeb, The Spatial Periodicity of Grid Cells Is Not Sustained During Reduced Theta Oscillations, Science (80-). 332 (6029) (2011) 592–595, ISSN 0036-8075, doi:10.1126/science.1201685.

M. P. Brandon, J. Koenig, J. K. Leutgeb, S. Leutgeb, New and Distinct Hippocampal Place Codes Are Generated in a New Environment during Septal Inactivation, Neuron 82 (4) (2014) 789–796, ISSN 08966273, doi:10.1016/j.neuron.2014.04.013, URL https://linkinghub.elsevier.com/retrieve/pii/S0896627314003031.

L. de Almeida, M. Idiart, J. E. Lisman, The Input-Output Transformation of the Hippocampal Granule Cells: From Grid Cells to Place Fields, J. Neurosci. 29 (23) (2009a) 7504–7512, doi:10.1523/JNEUROSCI.6048-08.2009.

P. Byrne, S. Becker, N. Burgess, Remembering the past and imagining the future: A neural model of spatial memory and imagery, Psychol. Rev. 114 (2) (2007) 340–375.

A. Bicanski, N. Burgess, A neural-level model of spatial memory and imagery, Elife 7 (7052) (2018) e33752, ISSN 2050-084X, doi:10.7554/eLife.33752.

T. Iijima, M. P. Witter, M. Ichikawa, T. Tominaga, R. Kajiwara, G. Matsumoto, Entorhinal-Hippocampal Interactions Revealed by Real-Time Imaging, Science (80-.). 272 (5265) (1996) 1176–1179, ISSN 0036-8075, doi:10.1126/science.272.5265.1176, URL http://www.sciencemag.org/cgi/doi/10.1126/science.272.5265.1176.

A. Guanella, D. Kiper, P. Vershure, A model of grid cells based on a twisted torus topology, Int. J. Neural Syst. 17 (04) (2007) 231–240, ISSN 0129-0657, doi:10.1142/S0129065707001093.

Y. Burak, I. R. Fiete, Accurate path integration in continuous attractor network models of grid cells., PLoS Comput. Biol. 5 (2) (2009) e1000291, ISSN 1553-7358, doi:10.1371/journal.pcbi.1000291.

E. Oja, Simplified neuron model as a principal component analyzer., J. Math. Biol. 15 (3) (1982) 267–273.

L. Slomianka, I. Amrein, I. Knuesel, J. C. Sørensen, D. P. Wolfer, Hippocampal pyramidal cells: the reemergence of cortical lamination, Brain Struct. Funct. 216 (4) (2011) 301–317, ISSN 1863-2653, doi:10.1007/s00429-011-0322-0, URL http://link.springer.com/10.1007/s00429-011-0322-0.

L. de Almeida, M. Idiart, J. E. Lisman, A Second Function of Gamma Frequency Oscillations: An E%-Max Winner-Take-All Mechanism Selects Which Cells Fire, J. Neurosci. 29 (23) (2009b) 7497–7503, doi:10.1523/JNEUROSCI.6044-08.2009.

D. Aronov, D. W. Tank, Engagement of Neural Circuits Underlying 2D Spatial Navigation in a Rodent Virtual Reality System, Neuron 84 (2) (2014) 442– 456, ISSN 0896-6273, doi:10.1016/J.NEURON.2014.08.042.

J. B. Hales, M. I. Schlesiger, J. K. Leutgeb, L. R. Squire, S. Leutgeb, R. E. Clark, Medial Entorhinal Cortex Lesions Only Partially Disrupt Hippocampal Place Cells and Hippocampus-Dependent Place Memory, Cell Rep. 9 (3) (2014) 893–901, ISSN 2211-1247, doi:10.1016/J.CELREP.2014.10.009.

K. M. Gothard, W. E. Skaggs, B. L. McNaughton, Dynamics of mismatch correction in the hippocampal ensemble code for space: Interaction between path integration and environmental cues, J. Neurosci. 16 (24) (1996) 8027–8040.

A. Samsonovich, B. L. McNaughton, Path integration and cognitive mapping in a continuous attractor neural network model, J. Neurosci. 17 (15) (1997) 5900–5920.

W. E. Skaggs, B. L. McNaughton, Spatial Firing Properties of Hippocampal CA1 Populations in an Environment Containing Two Visually Identical Regions, J. Neurosci. 18 (20) (1998) 8455–8466, ISSN 0270-6474, doi: 10.1523/JNEUROSCI.18-20-08455.1998.

V. H. Brun, M. K. Otnaess, S. Molden, H.-A. Steffenbach, M. P. Witter, M.-B. Moser, M. E. I., Place Cells and Place Recognition Maintained by Direct Entorhinal-Hippocampal Circuitry, Science (80-.). 296 (5576) (2002) 2243–2246, ISSN 00368075, doi:10.1126/science.1071089.

P. A. Naber, F. H. Lopes da Silva, M. P. Witter, Reciprocal connections between the entorhinal cortex and hippocampal fields CA1 and the subiculum are in register with the projections from CA1 to the subiculum, Hippocampus 11 (2) (2001) 99–104, ISSN 1050-9631, doi:10.1002/hipo.1028.

F. Kloosterman, T. van Haeften, M. P. Witter, F. H. Lopes da Silva, Electrophysiological characterization of interlaminar entorhinal connections: an essential link for re-entrance in the hippocampal-entorhinal system, Eur. J. Neurosci. 18 (11) (2003) 3037–3052, ISSN 0953-816X, doi:10.1111/j.1460-9568.2003.03046.x.

K. Mizuseki, K. Diba, E. Pastalkova, G. Buzsáki, Hippocampal CA1 pyramidal cells form functionally distinct sublayers, Nat. Neurosci. 14 (9) (2011) 1174– 1181, ISSN 1097-6256, doi:10.1038/nn.2894.

A. V. Masurkar, K. V. Srinivas, D. H. Brann, R. Warren, D. C. Lowes, S. A. Siegelbaum, Medial and Lateral Entorhinal Cortex Differentially Excite Deep versus Superficial CA1 Pyramidal Neurons, Cell Rep. 18 (1) (2017) 148–160, ISSN 2211-1247, doi:10.1016/J.CELREP.2016.12.012.

D. Mao, S. Kandler, B. L. McNaughton, V. Bonin, Sparse orthogonal population representation of spatial context in the retrosplenial cortex, Nat. Commun. 8 (1) (2017) 243, ISSN 2041-1723, doi:10.1038/s41467-017-00180-9.

W. E. Skaggs, B. L. McNaughton, Replay of neuronal firing sequences in rat hippocampus during sleep following spatial experience, Science (80-.). 271 (1996) 1870–1873.

G. Girardeau, K. Benchenane, S. I. Wiener, G. Buzsáki, M. B. Zugaro, Selective suppression of hippocampal ripples impairs spatial memory, Nat. Neurosci. 12 (10) (2009) 1222–1223, ISSN 1546-1726, doi:10.1038/nn.2384.

K. Benchenane, A. Peyrache, M. Khamassi, P. L. Tierney, Y. Gioanni, F. P. Battaglia, S. I. Wiener, Coherent theta oscillations and reorganization of spike timing in the hippocampal-prefrontal network upon learning, Neuron 66 (6) (2010) 921–936, ISSN 08966273, doi:10.1016/j.neuron.2010.05.013, URL http://linkinghub.elsevier.com/retrieve/pii/S0896627310003818.

H. Tanila, Hippocampal place cells can develop distinct representations of two visually identical environments, Hippocampus 9 (3) (1999) 235–246, doi: 10.1002/(SICI)1098-1063(1999)9:3%21235::AID-HIPO4%BF3.0.CO;2-3.

M. C. Fuhs, S. R. VanRhoads, A. E. Casale, B. McNaughton, D. S. Touretzky, Influence of Path Integration Versus Environmental Orientation on Place Cell Remapping Between Visually Identical Environments, J. Neurophysiol. 94 (4) (2005) 2603–2616, ISSN 0022-3077, doi:10.1152/jn.00132.2005, URL http://www.physiology.org/doi/10.1152/jn.00132.2005.

V. Paz-Villagrán, E. Save, B. Poucet, Spatial discrimination of visually similar environments by hippocampal place cells in the presence of remote recalibrating landmarks, Eur. J. Neurosci. 23 (1) (2006) 187–195, ISSN 0953816X, doi:10.1111/j.1460-9568.2005.04541.x, URL http://doi.wiley.com/10.1111/j.1460-9568.2005.04541.x.

H. J. Spiers, R. M. A. Hayman, A. Jovalekic, E. Marozzi, K. J. Jeffery, Place Field Repetition and Purely Local Remapping in a Multicompartment Environment, Cereb. Cortex 25 (1) (2015) 10–25, ISSN 1047-3211, doi: 10.1093/cercor/bht198.

R. M. Grieves, B. W. Jenkins, B. C. Harland, E. R. Wood, P. A. Dudchenko, Place field repetition and spatial learning in a multicompartment environment, Hippocampus 26 (1) (2016) 118–134, ISSN 10509631, doi: 10.1002/hipo.22496, URL http://doi.wiley.com/10.1002/hipo.22496.

P.-Y. Jacob, G. Casali, L. Spieser, H. Page, D. Overington, K. Jeffery, An independent, landmark-dominated head-direction signal in dysgranular retrosplenial cortex, Nat. Neurosci. 20 (2) (2017) 173–175, ISSN 1097-6256, doi: 10.1038/nn.4465, URL http://www.nature.com/articles/nn.4465.

B. L. McNaughton, C. A. Barnes, J. L. Gerrard, K. Gothard, M. W. Jung, J. J. Knierim, H. Kudrimoti, Y. Qin, W. E. Skaggs, M. Suster, K. L. Weaver, Deciphering the hippocampal polyglot: the hippocampus as a path integration system, J. Exp. Biol. 199 (Pt 1) (1996) 173–185, ISSN 0022-0949.

E. S. Rosenzweig, A. D. Redish, B. L. McNaughton, C. A. Barnes, Hippocampal map realignment and spatial learning., Nat. Neurosci. 6 (6) (2003) 609–15.

